# Simultaneous cortical tracking of competing speech streams during attention switching

**DOI:** 10.1101/2025.07.02.662762

**Authors:** Sara Carta, Emina Aličković, Johannes Zaar, Alejandro López Valdés, Giovanni M. Di Liberto

**Author notes:** Senior author (Alejandro López Valdés, Giovanni M. Di Liberto).

## Abstract

Successful speech communication in multi-talker scenarios requires a skilful combination of sustained attention and rapid attention switching. While the neurophysiology literature offers detailed insights into the neural underpinnings of sustained attention, there remains considerable uncertainty on how attention switching takes place. In this study, using EEG recordings from normal-hearing adults in an immersive multi-talker environment, we measured the neural encoding of two competing speech streams amid background babble. Participants were cued to switch attention between streams every 15–30 seconds. Neural tracking was assessed via Temporal Response Functions (TRF), confirming reliable decoding of attentional focus. Our results indicate asymmetric disengagement and engagement processes during attention switches, where the neural tracking of the new target stream emerges before disengaging from the previous target, revealing a transient simultaneous encoding of two speech streams. That transition was closely mirrored by a reduction in EEG alpha power, informing on the cognitive effort during different phases of the attention switch. We then isolated cortical activity reflecting lexical prediction mechanisms to determine how lexical context is updated after an attention switch, comparing four numerical hypotheses that were constructed using Large Language Models. Our findings elucidate both the temporal and contextual mechanisms underlying auditory attention shifts, pointing to the possibility that listeners carry out a reset in lexical context after switching attention. By focusing on dynamic attentional reallocation, this study offers insights into the brain’s capacity for flexible speech processing in complex listening environments.

## Introduction

To understand speech in multi-talker environments, listeners single out the target speaker from competing sound streams ^1–3^. The neurophysiology of this selective attention process has been widely studied with simulated *cocktail-party* scenarios ^4,5^, shedding light on how our brains segregate a target stream from competing speech streams, and enabling the transformation of the target speech into linguistic meaning. While the extent to which masker speech streams are processed remains highly debated ^6–8^, there is no doubt that there are considerable differences between the processing of target and masker speech, which have been measured with various technologies, such as non-invasive electroencephalography, EEG ^1,9^, intra-cranial electroencephalography, iEEG ^10^, magnetoencephalography, MEG ^3,11^ and functional magnetic resonance imaging, fMRI ^12,13^. That work could pinpoint precise loci in the auditory cortical areas where that segregation emerges ^14^ as well as measuring the substantial (but not total) suppression of linguistic processing for the masker speech ^1,15–17^. However, neurophysiology literature in this field has almost entirely focussed on sustained attention tasks ^2,10^, leaving considerable uncertainty on the neural underpinnings of attention switching.

Dynamic switching paradigms have been widely used in the domain of cognitive control studies to probe for cognitive flexibility and cognitive stability ^18^. In those experiments, participants are often required to flexibly adapt their behavioural response depending on new instructions, initiating a task-switch ^19–21^. For example, given a single digit, they are required to classify it either based on parity, i.e. whether it is even or odd, or based on relative magnitude, i.e. whether the digit is greater than or less than 5 ^22^. In these paradigms, the switch-cost is the increase in reaction time or error rate when switching from one task to the other. Similar behavioural paradigms have also involved simple speech stimuli in multi-talker settings ^23–25^. However, the main interest of those tightly controlled experiments was to model the process of target speech selection as one particular instance of a task-switching problem, i.e. target stream selection could either depend on spatial location or voice identity ^23^, rather than focussing on the dynamic aspect of attention re-allocation per se in naturalistic multi-talker scenarios. As such, very little is known on how a flexible reorienting of attention might impact speech processing of continuous competing streams.

In recent speech neurophysiology research, experimental paradigms have started to include switches of attention as a tool towards tailored EEG/MEG methodological advances in the domain of attention decoding ^26,27^, or to investigate how sustained speech attention unfolds for moving auditory objects ^28^. However, to the best of our knowledge, only one previous study has specifically focussed on the neurophysiology of attention switching in multi-talker scenarios, relating the neural encoding of speech during attentional re-orienting with EEG alpha activity and pupil dilation dynamics ^29^. Those findings proved that the neurophysiology of attention switching can be studied non-invasively. Building on that work, our study sheds light on the exact neural dynamics supporting the steering of attention between two competing speech streams, disengaging from the previous target stream while engaging to the new one.

In this study, we measure the neural encoding of speech at millisecond resolution as listeners steer their attention from one speaker to another. We test whether engagement with a new speech stream begins before before disengagement from the previous target is complete, resulting in a brief period of simultaneous tracking of both streams. Such an asymmetry in the disengagement-engagement processes, even if transient, could support the ability to explore alternative auditory streams while maintaining attention to a given stream ^30^.

The neural encoding of speech was measured from normal-hearing adult participants using EEG during an immersive multi-talker listening task. Participants were exposed to two competing speech streams from TED talks, presented via two front-facing loudspeakers, while background noise from a 16-talker speech babble played from rear loudspeakers (**Figure 1A**). An on-screen arrow cued participants to attend to one of the two speech streams and to shift their attention rapidly whenever the arrow changed direction, approximately every 10–30 seconds (**Figure 1B**). Neural tracking of target and masker speech was quantified using the Temporal Response Function (TRF), describing the linear relationship between each speech stream and the neural responses. As an initial validation, we confirmed that the attended stream could be reliably decoded from the EEG, consistent with the extensive literature on sustained attention ^9,10,31^. This confirms that the EEG responses in this experiment reflects differential encoding of target versus masker speech (**Figure 1C**).

**Figure 1.**
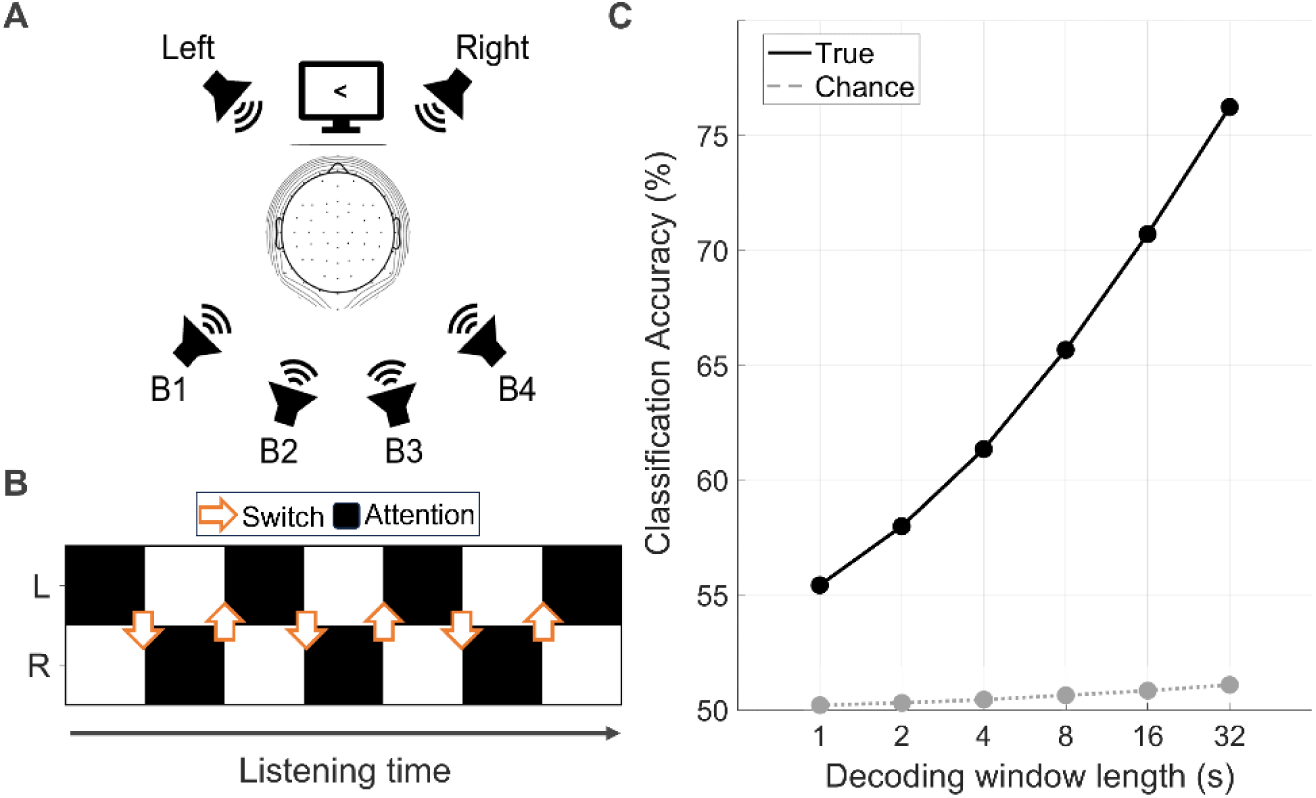
Experiment overview and validation. **(A)** Participants were presented with speech from two loudspeakers placed in front of them with 60° of separation (30° left and 30° right), and with concurrent 16-talker background noise (B1-B4). In each trial, the screen presented an arrow pointing to the target speech stream. Participants were instructed to switch attention as soon as the visual cue changes direction. **(B)** Schematic diagram of one experimental trial. The black area represents blocks of attention either to the left (L) or right (R) front streams. The red arrows indicate the instants where the attention cue switches side (six times per trial). Note that block duration was randomised and always between 15 and 30 seconds, with trials lasting 3 minutes. **(C)** EEG data validation was carried out by running an attention decoding analysis. Progressively longer decoding windows were considered (larger windows use more data, typically leading to more accurate decoding scores). Binary classification scores are reported arbitrating between the target and masker streams. The dashed line indicates the 95^th^ percentile of a random distribution calculated by randomising the classification labels. Statistically significant attention decoding classification scores were measured for all the decoding windows considered, with numerical results comparable with previous studies on selective attention ^31–33^.

We next addressed two fundamental questions about the neural mechanisms underlying attention switching in naturalistic listening. First, we asked whether the processes of engaging with a new speech stream and disengaging from a previous one unfold symmetrically (**Figure 2 and 3**). To test this, we fit encoding TRF models to EEG data, measuring the neural tracking of the two competing speech streams over time. This allowed us to characterise the average encoding dynamics surrounding attention switches, comparing disengagement and engagement processes. The second objective was to understand how our brains update and use lexical context when switching attention (**Figure 4**). We formulated four hypotheses for this phenomenon and built quantitative predictions for each using a state-of-the-art large language model (LLM). This resulted in four TRF regressors reflecting lexical surprise with a different sensitivity to previous context and switch occurrence, for the different hypotheses. Encoding TRF models were fit for the four hypotheses, revealing the most physiologically plausible lexical processing mechanism.

**Figure 2.**
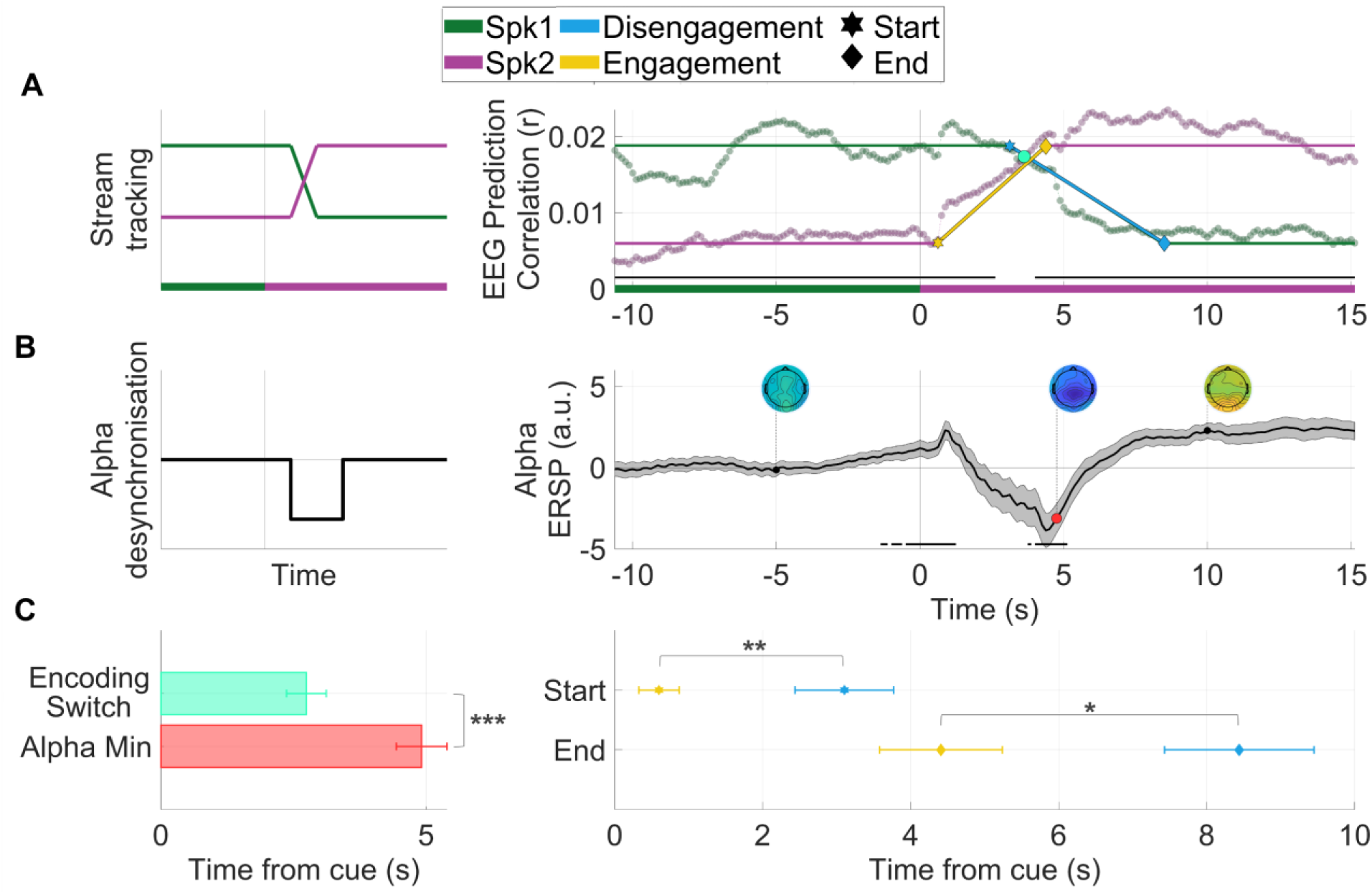
The attention-switching cue prompts a robust disengagement from Speaker 1 and engagement to Speaker 2, and it is followed by a significant decrease in the EEG alpha ERSP. Disengagement has longer temporal dynamics compared to engagement. (A) Left: Speech tracking encoding for an attention switch from Speaker 1 and 2. The trajectory in the panel represents our null hypothesis, where the disengagement and engagement processes progress in a symmetric manner after the switch-cue (vertical grey line). **Right:** Results for the neural tracking of Speaker 1 and Speaker 2 across the switching cue. EEG prediction correlations (average across all channels) obtained from a 4-s sliding-window TRF model including Envelope (Env), Word Onset (WO) and Word Surprisal (WS) features. Coloured horizontal bars at the bottom of the plot indicate the attention instruction around the attention switching cue. The turquoise dot indicates the encoding switch of EEG prediction correlations based on Spk1- and Spk2-speech features. The piecewise linear model fit for disengagement and engagement is overlayed on the EEG prediction correlation values (average of all channels and participants). Hexagram shapes indicate the start of the disengagement (blue) and engagement (yellow) processes, while diamonds represent the end of the transitions. **(B) Left:** Diagram of expected results for alpha-band ERSP (event-related spectral perturbation) across the switching cue. **Right:** ERSP of the alpha band (8-12 Hz) around the switching cue (average of all channels), computed with a 4-s sliding-window, as above. Scalp topographies at selected time points reveal a pattern of posterior negativity, which drops significantly following the instruction to switch (thick black lines indicate a statistically significant change compared to pre-switch baseline). The red dot represents the average of ERSP minima across participants. The shaded area represents the standard error of the mean (SEM) across participants. **(C) Left:** Comparison of encoding switch of EEG prediction correlations (turquoise bar) and alpha ERSP minimum (red bar) for a 4-s sliding-window. The alpha ERSP reaches its minimum significantly after the Spk1-Spk2 encoding switch point. **Right:** Comparison of temporal dynamics for start and end points of disengagement and engagement processes. Stars indicate significant statistical effects (paired sample t-tests; *p ≤ 0.05; **p ≤ 0.01; **p ≤ 0.001).

**Figure 3.**
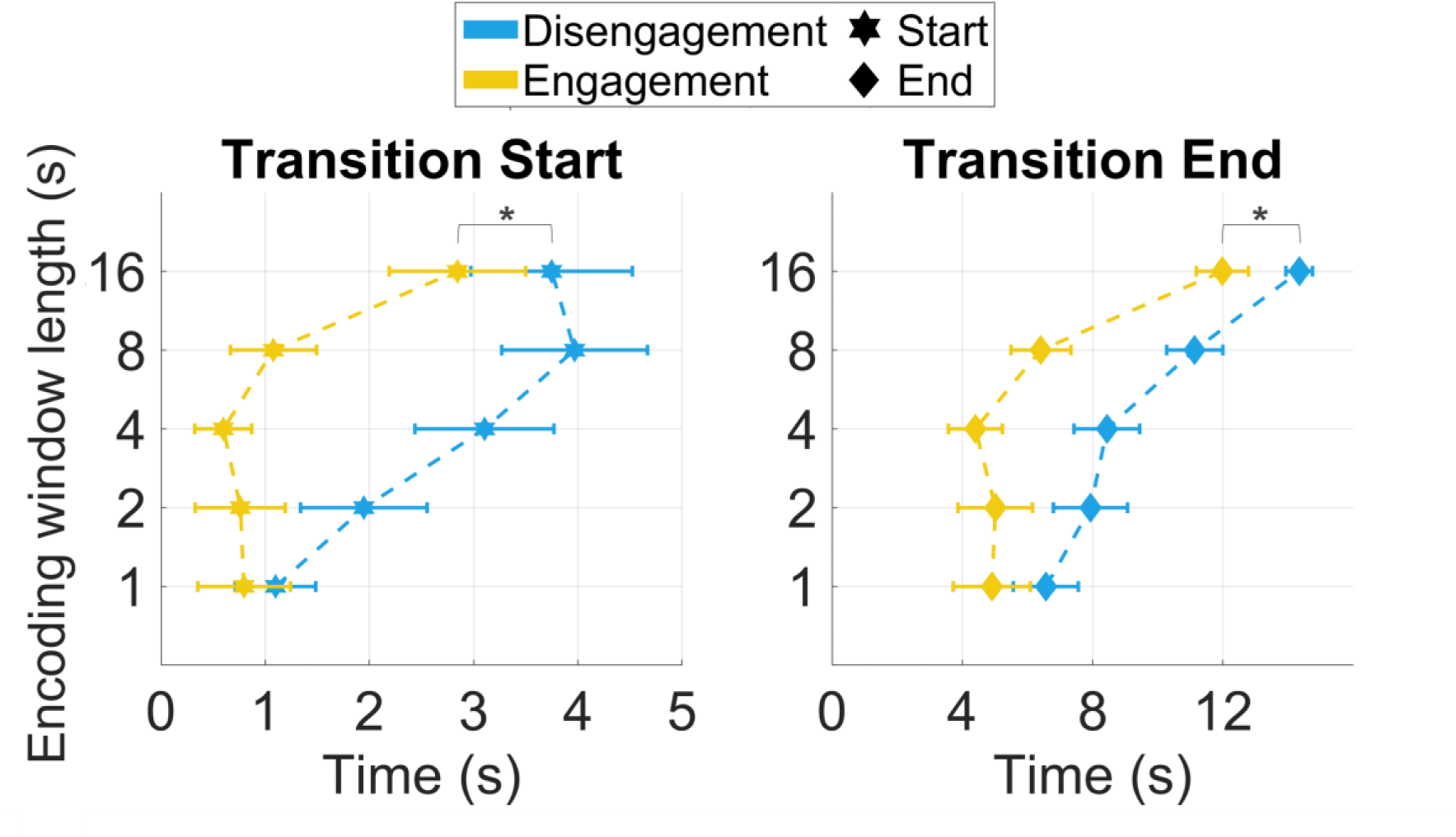
Comparing the start and end transition points for the disengagement and engagement processes after the attentions switching cue. The process of engaging to a new speaker begins and ends significantly earlier than disengaging from the previously attended speaker. (A-B) Start and end points of the transition for the disengagement (blue) and engagement (yellow) processes over five TRF sliding window lengths. Error bars represent SEM across participants. Stars indicate significant effects of process type (two-way repeated measures ANOVA; *p ≤ 0.05; **p ≤ 0.01; ***p ≤ 0.001).

**Figure 4.**
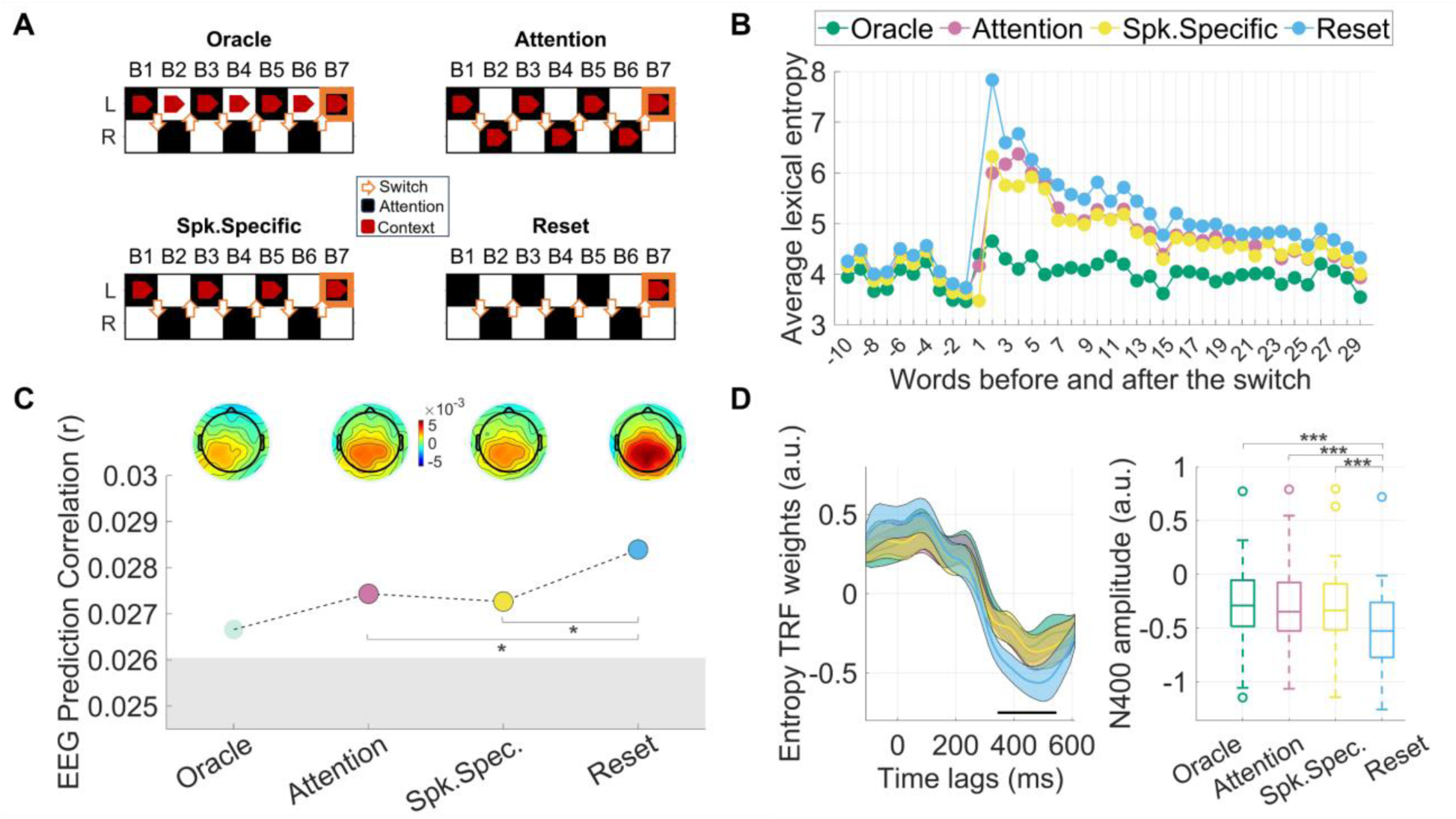
Investigating lexical prediction mechanisms during attention switching. **(A)** Layout of the four context models. Blocks coloured in black illustrate sustained attention either to the Left or Right stream, while orange arrows indicate attention switching cues. The thick red arrow indicates the context used to guide word predictions for the current block (B7, highlighted in orange). **(B)** Average lexical entropy at words preceding and following the attention switch cue. Note that no value for entropy is displayed in the Reset model for the first word after the switch, due to the context being fully reinstated. **(C)** EEG prediction correlations for the four multivariate TRF models, only differing in their entropy feature. Coloured dots indicate the average across all electrodes and participants. The grey area at the bottom represents the average encoding accuracy of a multivariate TRF without any semantic information (Envelope + Word Onset). Stars represent statistically significantly greater EEG prediction correlations for the Reset model compared to the other models (Significance levels: *p < 0.05, **p < 0.01, ***p < 0.001). Topographical patterns illustrate the gain due to semantic information (compared to the Envelope + Word Onset TRF) for the four models. **(D)** (Left) TRF weights for the entropy feature at time-lags between -100ms and 600ms relative to stimulus onset. Transparent shaded areas represent the standard error of the mean (SEM) across participants. The horizontal black line indicates the time window employed to compute the average TRF-N400 amplitude. (Right) Boxplots representing the distribution of the TRF-N400 amplitude across participants for the four context models. The central line within each box represents the median, while the edges of the box indicate the interquartile range (IQR). Whiskers extend to the most extreme data points within 1.5 times the IQR from the quartiles. Outliers are plotted as individual points beyond the whiskers. Stars indicate statistically significant comparisons (Significance levels: *p < 0.05, **p < 0.01, ***p < 0.001).

This study provides substantial new insights into the temporal unfolding and contextual mechanisms guiding attention switching, encompassing both low and high levels of speech abstraction.

## Results

### Behavioural performance

Following each trial, participants were presented with a four-alternative forced-choice question about the content of one of the speech streams to confirm task engagement. Behavioural performance revealed that participants were able to successfully reply to comprehension questions, with an average accuracy of 86.3% (SEM 2.6%). Due to technical issues, behavioural data for one of the twenty-four participants was not available, therefore behavioural performance was computed based on the data from the remaining twenty-three participants.

### Decoding of selective auditory attention in a dynamic switching scenario

Participants’ attention was decoded with a backward TRF analysis, describing the relationship between the EEG signals and the envelope of the target speech. For each left-out trial, the speech envelope reconstructed from the target decoding model was correlated with the envelopes of both the left and right speech streams. Attention was classified by determining which speech stream’s envelope showed a higher correlation with the reconstructed envelope. Since this was a dynamic attention-switching scenario, the attended speech could alternatively correspond to the left or the right stream. Classification was considered correct when the reconstructed envelope correlated more strongly with the target speech envelope than with the masker envelope. Classification accuracy was then computed as the proportion of instances where this criterion was met. To establish chance performance, left and right labels were randomly shuffled 100 times for each decoding window. As shown in **Figure 1C**, the longer decoding windows led to higher classification performances. However, even with a 1-second window, classification accuracy was significantly above chance level, and all decoding windows yielded classification rates significantly above the 95^th^ percentile of its chance distribution (paired two-tailed t-test, FDR-corrected for multiple comparisons for windows of 1s, 2s, 4s, 8s, 16s, 32s, respectively: *p* = 0.47e-9; 0.53e-9; 0.27e-9; 0.24e-9; 0.24e-9; 0.24e-9). These findings align with previous work decoding sustained attention or employing match-vs-mismatch classification metrics ^9,31,34^, and confirm that a classification based on the envelope reconstruction can reliably track selective attention even during attention switches.

### Neural tracking of competing speech streams in a dynamic switching scenario reflects the listener’s focus of attention and is related to changes in alpha ERSP

A multivariate TRF analysis was carried out to characterise the neural tracking of two competing speech streams in a setting where participants were instructed to dynamically switch their attention between the two streams. Single-subject TRFs were trained on the target stream and tested on both speakers (i.e., Spk1 and SpK2) using a multivariate speech representation that included Envelope, Word Onset and Word Surprisal features (for more details see Methods). EEG prediction correlations were computed using a sliding window to correlate true and predicted EEG signals over time, with a leave-one-out cross validation procedure, and were averaged across all EEG channels. In order to analyse robust switching dynamics, we selected 19 participants displaying a reliable attentional bias over the course of the switch, based on an above-chance classification accuracy criterion (>50%) over the course of the switch. Aligning with our expectation (**Figure 2A**), EEG prediction correlations around the switching cue reflected tracking of Spk1 and Spk2 streams consistent with the attention instructions, such that Spk1 was significantly more tracked than Spk2 before the switch, while the reverse pattern was observed after the switch (paired two-tailed t-test of Spk1-Spk2 difference against zero, FDR-corrected for multiple comparisons, *p* < 0.005).

As the attention switch unfolds, also the grand-mean ERSP in the alpha frequency band displayed a statistically significant change compared to baseline (one-sample t-test against zero, FDR-corrected for multiple comparisons), revealing a pattern of occipito-parietal negativity in the scalp topographies (**Figure 2B**). This is consistent with our expectation of an impact of attentional reorientation on the EEG alpha band, which has already been shown to reflect attention switching behaviour in competing speech listening scenarios ^29^.

EEG prediction correlations for Spk1 and Spk2 converged, before significantly separating again once the switching process was concluded and, presumably, the attention was fully reallocated. Here, we refer to the time point when EEG prediction overlap between Spk1 and Spk2 as the encoding switch point. Given the observed statistically significant drop in alpha ERSP, we asked how the temporal dynamics of this drop compared to those of the EEG prediction correlations. To address this, for each participant, we identified the time of the alpha ERSP minimum, and the encoding switch point, based on an encoding window of 4s (**Figure 2C**). The choice of this particular encoding window for our main analysis is justified based on the classification accuracy results (**Figure 1C**), since it is a good compromise between temporal resolution and classification performance. However, the same pattern of results holds when considering multiple encoding windows simultaneously (**Figure S1**). A paired t-test comparing the temporal dynamics of the alpha ERSP and the EEG prediction correlations showed that the minimum of the alpha ERSP drop significantly follows the encoding switch point (*t*(18) = 3.85, *p* = 0.01, Cohen’s *d* = 0.88).

We then evaluated multiple encoding window lengths, assessing the effect of Metric (encoding switch vs. ERSP minimum) and Window (1s, 2s, 4s, 8s) on the timing of the encoding switch and the minimum of the alpha ERSP with a 2-way repeated measures ANOVA. The analysis revealed that the temporal dynamics of both encoding switch and alpha ERSP minimum became longer as the encoding window length increased (*F*(1,18) = 27.87, *p* = 8.77e-8, η_p_^2^ = 0.61; a Greenhouse-Geisser’s correction applied due to sphericity violation), which is unsurprising given the methodological constraints we discussed (see **Methods**). More interestingly for our question, a statistically significant effect of Metric emerged (*F*(1,18) = 7.91, *p* = 0.012, η_p_^2^ = 0.3), with the alpha ERSP minimum occurring significantly later than the encoding switch across a range of encoding windows (Holm-corrected post-hoc *t*-test: *t*(18) = 2.81, *p* = 0.012, Cohen’s *d* = 0.65).

### Dissecting the temporal dynamics of attentional disengagement and engagement during attention switching

The attention switching cue prompts the listener to reallocate their attention from the previously attended speaker, Spk1, to the newly attended speaker, Spk2. While this re-routing of attention appears to be a single, unified process, it is possible to distinguish two separate operations that are necessary for it to happen: disengagement, which we define as the decrease in neural tracking for the previously attended speech stream, and engagement, which we define as the increase in neural tracking for the previously unattended speech stream. Our goal was to clarify the temporal dynamics of these two operations to understand whether they occur fully in parallel, serially, or with a certain degree of overlap. As above, we first selected participants displaying a reliable attentional bias over the course of the switch (see **Methods**). For this selection of 19 participants, we fitted a piecewise linear regression on single-subject EEG prediction correlations, and found the optimal breakpoints, corresponding to the start and end time points of disengagement and engagement (**Figure 2C**). As above, we chose to focus on an example window of 4s and later replicated our results on a range of encoding window lengths. Disengagement and engagement processes were compared separately based on their start times and end times, revealing consistently earlier temporal dynamics for the engagement compared to the disengagement.

Engagement to the newly attended speaker started significantly earlier than the disengagement from the previously attended speaker (paired-sample *t*-test: *t*(18) = 3.06, *p* =0.007, Cohen’s *d* = 0.43), and finished significantly earlier (paired-sample *t*-test: *t*(18) = 2.56, *p* = 0.02, Cohen’s *d* = 0.42).

We then extended our analysis to a range of sliding window lengths and compared start times and end times for disengagement and engagement processes, including Window (1s,2s,4s,8s,16s) and Process (disengagement vs. engagement) as main factors in a repeated measures ANOVA. Regarding the start points (**Figure 3A**), our analyses revealed an expected statistically significant effect of Window (*F*(1,18) = 8.37, *p* = 0.003, η_p_^2^ = 0.32; the assumption of sphericity was not met; hence, a Greenhouse-Geisser’s correction was applied), with longer temporal dynamics corresponding to longer encoding window lengths. More importantly, we also observed a significant main effect of Process (*F*(1,18) = 7.22, *p* = 0.015, η_p_^2^ = 0.27), with engagement to the newly attended stream starting significantly earlier than the disengagement to the previously attended stream (Holm-corrected post-hoc *t*-test: *t*(18) = 2.69, *p* = 0.015, Cohen’s *d* = 0.64). The same statistical analysis was repeated separately on the end time points of disengagement and engagement processes (**Figure 3B**), revealing once again a main effect of Window, whereby longer encoding windows yield longer temporal transitions (*F*(1,18) = 36.7, *p* = 6.8e-8, η_p_^2^ = 0.32; the assumption of sphericity was not met; hence, a Greenhouse-Geisser’s correction was applied). A significant main effect of Process also emerged (*F*(1,18) = 6.36, *p* = 0.021, η_p_^2^ = 0.11), revealing that the process of engagement to the newly attended speaker, not only starts, but also ends significantly earlier than the disengagement (Holm-corrected post-hoc *t*-test: *t*(18) = 2.52, *p* = 0.021, Cohen’s *d* = 0.75).

### Determining how lexical predictions are built during attention switching

Reorienting attention to a different speech stream implies a change of context and, consequently, different semantic priors for lexical predictions. We thus hypothesised that incorporating this change of context into the structure of our semantic regressor in a multivariate encoding TRF model would increase EEG prediction correlations, as it would better reflect the dynamically updating neural tracking of the competing speech streams. We compared four alternative models representing how context could be incrementally accumulated for performing lexical predictions at one particular attention block (e.g., B7, in **Figure 4A**). A naïve Oracle model, which uses all available context of previous blocks from the current stream —whether attended or unattended— to predict words from the current block, served as our baseline, since it was essentially a switch-unaware contextual representation. Speaker-Specific and Attention models were instead switch-aware models, as they only considered previously attended blocks as part of the context for lexical predictions. Speaker-Specific assumed a higher degree of stream segregation, since its context only consisted of previously attended blocks from the same speech stream, while Attention included any previously attended block from both streams. The Reset model instead ignored all previously attended blocks from any of the streams and computed context only over the course of the current block of attention, as if the priors for lexical predictions were reset at each attention switch (**Figure 4A**).

As lexical entropy is a proxy of uncertainty for next-word prediction, its values should be impacted by a switching cue, which determines an abrupt change of context. **Figure 4B** shows the change of average lexical entropy values in words preceding and following the switch cue, which vary depending on the context models. It can be observed that the Reset model peaks with the highest uncertainty and slowly decays over the course of the next words, while the Attention and Speaker Specific models have overall similar lexical entropy dynamics and more stable values. Consistently with its switch-unaware nature, the Oracle model instead displays entropy values that are largely unchanged despite the switch.

Lexical surprisal and lexical entropy were used as semantic information regressors for each context model and separately included in a multivariate stimulus representation to fit single-subject encoding TRFs (Envelope-Word Onset-Word Surprisal and Envelope-Word Onset-Word Entropy). Resulting TRF weights and EEG prediction correlations were then compared across context models, with the hypothesis that switch-aware and context-rich representations (e.g., Speaker-Specific or Attention) would best describe neural activity in attention-switching scenarios.

Before comparing the context models, we first tested whether each of them yielded a significant encoding accuracy gain compared to the baseline model only consisting of acoustic features (Envelope and Word Onset). When using entropy as a regressor for semantics, all models, with the exception of Oracle, showed a statistically significant gain, suggesting a robust tracking of semantic information in addition to the stimulus acoustics (paired t-tests: Oracle vs. Acoustics: *p* = 0.29; Spk.Spec. vs. Acoustics: *p* = 0.03; Attention vs. Acoustics: *p* = 0.03; Reset vs. Acoustics: *p* = 0.003). Employing word surprisal as semantic regressor yielded similar results, with all the models showing a robust encoding of semantic information, apart from Oracle (paired t-tests: Oracle vs. Acoustics: *p* = 0.16; Spk.Spec. vs. Acoustics: *p* = 0.04; Attention vs. Acoustics: *p* = 0.03; Reset vs. Acoustics: *p* = 0.02). The non-significant gain of the Oracle model compared to the acoustic model was expected, since Oracle was designed as a control *switch-unaware* model.

In contrast to our expectation, the Reset context model was shown to yield higher EEG prediction correlation values when entropy was used as a regressor for semantics (**Figure 4C**). A repeated measures ANOVA revealed a statistically significant effect of the main factor, Context Model (*F*(1,23) = 7.12, *p* = 0.001, η_p_^2^ = 0.27; with Greenhouse-Geisser’s correction). In the Holm-corrected post-hoc tests, the Reset model was shown to yield significantly higher encoding accuracies than Oracle (*t*(23) = 4.55, *p* = 1.32e-4, Cohen’s *d* = 0.14), and Speaker Specific (*t*(23) = 2.96, *p* = 0.021, Cohen’s *d* = 0.09). Reset was also marginally significant compared to the Attention model (*t*(23) = 2.52, *p* = 0.056, Cohen’s *d* = 0.08). We then assessed the difference of TRF weights for the entropy feature across the four context models (**Figure 4D**), averaging the weights’ amplitude within a window broadly centred around the TRF-N400 latency (350ms-550ms). A repeated measure ANOVA was run on the weights’ amplitude values, revealing a main effect of Context Model (*F*(1,23) = 14.56, *p* = 1.9e-7, η_p_^2^ = 0.39; a Greenhouse-Geisser’s correction was applied). Post-hoc tests (Holm-corrected) showed that weights for the Reset model had lower TRF-N400 amplitude compared to Oracle (*t*(23) = -5.93, *p* = 6.41e-7, Cohen’s *d* = 0.56), Attention (*t*(23) = -4.94, *p* = 2.1e-5, Cohen’s *d* = 0.47), and Speaker Specific (*t*(23) = -5.11, *p* = 1.4e-5, Cohen’s *d* = 0.48).

When fitting a multivariate TRF including lexical surprisal as a semantic regressor we were not able to observe a statistically significant difference between the four context models for the EEG prediction correlations, even if the Reset model yielded overall higher accuracy values. No difference was observed also when comparing the TRF-N400 amplitude of the models’ TRF weights.

## Discussion

Speech communication in multi-talker environments requires a skilful combination of sustained attention and rapid attention switching abilities ^5,30^. While the neurophysiology of sustained speech attention has been widely studied ^1,9,35,36^, less is known about the neural mechanisms of attention switching. Here, we fill this gap with a tailored EEG experiment examining the neurophysiology of attention switching across different levels of speech abstraction. In doing so, we 1) demonstrated an experimental paradigm that can successfully probe both sustained attention and attention switching mechanisms; 2) successfully dissected disengagement and engagement processes with a high temporal resolution, identifying substantial asymmetries in their temporal unfolding and a transient simultaneous encoding of two speech streams; and 3) proposed a neurophysiologically plausible explanation of how our brains update and use lexical context when switching attention.

The findings in this study have several implications for our understanding of speech attention switching mechanisms. The asymmetry measured between disengagement and engagement processes highlights the importance of studying the two processes separately. That distinction was often not considered in previous studies on sustained attention, which often focussed on measures of attention bias or classification ^10,31,32,37^. The effectiveness of such decoding metrics has been a driving force for research on brain-computer interfaces such as cognitively-controlled hearing devices ^38–41^. Our finding highlights that encoding metrics enable a sufficient level of detail for disentangling how the encoding of different streams evolves over time. Here, we measured an asymmetry between disengagement and engagement processes during attention switching in a very specific scenario. Indeed, it will be important to determine how that relationship is modulated by factors such as cognitive load, ageing, cognitive abilities, hearing difficulties, interest in the speech content, frequency of attention switches in a trial, among many others. Of course, future work should also scrutinise how the specific nature of the task might impact that phenomenon.

Sustained attention tasks, where participants focus on a target speech while ignoring the masker ^2,42^, involve a quite particular scenario where listeners have no incentive to monitor unattended streams. In real-life situations, however, listeners may have reasons to explore alternative speech streams, for example due to a lack of interest in the current speaker. Our experimental paradigm more closely mirrors this scenario. Although the instructed nature of the task makes it less realistic, the paradigm incentivises monitoring the masker and being ready to rapidly switch attention, which contrasts with sustained attention tasks. While the asymmetry observed may be specific to this experimental paradigm, the result indicates that our brains can engage with a new target even before starting the disengagement from the previous one. This *de facto* produces a transient simultaneous cortical tracking of two speech streams, with both measurements comparable to sustained attention in strength, aligning with auditory scene monitoring mechanisms ^43,44^. Intuitively, maintaining a transient parallel representation of multiple speech sources during attention switching is an efficient neural processing strategy. It allows the flexibility to switch back to the previous stream, if necessary, without fully committing to the newly attended speech immediately. This phenomenon supports previous claims that our brains can process speech maskers beyond the acoustic level, encoding linguistic properties to some extent ^7,8,16^.

Unattended speech streams are also represented in human cortical activity, with evidence for lower encoding strengths or longer time latencies than the target stream ^1,45,46^, and with a gradient of attentional bias from primary to nonprimary auditory cortex ^13,47^, whereby the unattended stream encoding tends to be substantially reduced or not measurable in higher-order cortical areas ^48,49^. Prior research has shown that not only the speech envelope, but also other key features of the unattended speech, such as acoustic onsets, are neurally represented ^50,51^ and, when not readily available due to speech masking, they are even restored at later temporal scales ^52^. One interpretation is that encoding a template structure of the unattended speech might be a useful strategy to suppress it ^50^ . Other research found attentional fluctuations corresponding to the changes in Target-Masker relative sound energy, with evidence that our brains may encode some phonetic information of unattended streams ^53,54^. The encoding of such unattended stream information may be one of the factors facilitating the rapid engagement during attention switching.

Using alpha ERSP as indicators of listening effort, this study related perceptual demands with attention switching dynamics. A large reduction in EEG alpha-band power was measured consistently about 4.5 seconds after the attention-switching cue. The ERSP trajectory suggests that a strong listening effort persists throughout the attention switch, with a substantial reduction close to the end of the switch (**Figure 2A**). The late ERSP trough—marking a turning point in listening effort—occurred significantly later than the moment of the attention switch itself. This finding extends prior research on the neural correlates of selective attention and listening effort. Variations in alpha ERSP have been associated not only with auditory attention effort ^55–59^, but also with the active suppression of irrelevant information in accordance with behavioral goals ^60–62^. Notably, cognitive load and the inhibition of irrelevant stimuli appear to be even more strongly influenced by attention reorienting than by maintaining attention on a single speaker ^29,63^, highlighting the increased cognitive demands of switching attention. While alpha power has commonly been linked to subjective listening fatigue ^55,57^ or the signal-to-noise ratio (SNR) between attended and unattended streams ^58^, it is less frequently associated with neural markers of speech tracking ^64,65^. In this study, we identified a link between the temporal dynamics of neural speech tracking and listening effort, suggesting a potentially valuable metric for future research into attention-switching challenges.

This study examined how linguistic context is dynamically updated during attention switching by comparing four context-accumulation models that varied in their sensitivity to attention switches and access to prior context: an Oracle model (context-rich but switch-unaware), Speaker Specific and Attention models (both context-rich and switch-aware but differing in stream selectivity), and a Reset model (aware of the switch but limited to the current block’s context). For each model, a multivariate temporal response function was fit using a semantic regressor aligned with the respective context strategy. Unexpectedly, the Reset model best predicted EEG data, outperforming models that retained past context and challenging the assumption that prior semantic information aids comprehension during attention switches. This finding may reflect a cognitive mechanism, suggesting listeners reset context and recalibrate their lexical predictions dynamically when switching attention to a new stream. Another explanation highlights that predictive nature of LLMs like Mistral, which are optimised for next-word prediction, without a requirement of being neurophysiologically plausible. As such, despite consistent reports of the growing similarity between more modern LLMs with neurophysiological activity ^66–68^,the higher neurophysiological plausibility of the Reset model tells us that prior context is not availed of by our brains in the way implemented by the competing models, Attention and Spk-Specific. Nonetheless, we don’t exclude that other strategies making use of the prior context might be in place. One possibility is that a dynamic attention switching task prompts a different use of that context, for example by capturing its essence in a more abstract and less specific way, as the gist of the story ^16^. As such, there could be value in exploring different strategies for context representation, for example by employing Large Concept Models ^69^, which are trained and optimised for sentence prediction. This latter possibility is also supported by recent work on the accumulation of linguistic context in LLMs and the human brain ^70^ while listening to continuous monologues, showing that LLMs with a limited context window (32 tokens) and with access to a coarse summary of the previous context predict neural activity better than LLMs with a higher token-memory.

In summary, this study showed that the process of attention switching in a realistic multi-talker scenario can be investigated in terms of its engagement and disengagement components, with a transient parallel representation of the two streams. We highlight the importance of relating metrics of neural tracking of speech with metrics of listening effort and demonstrate that the listening effort starts decreasing following successful disambiguation of the two streams during attentional re-allocation. Finally, we introduce an approach for modelling lexical context of dynamic attention scenarios, showing the sensitivity of transformer-based language models to subtle differences in context accumulation strategies. These findings have implications for future investigations into the cortical mechanisms of attention re-orienting and can be employed to highlight differences across diverse populations in terms of age and hearing levels.

## Methods

### Participants and experimental procedure

For the present study, which was approved by the Ethics Committee of the School of Psychology at Trinity College Dublin, we recruited 24 young native English speakers (between 18 and 39 years of age), who gave their written informed consent. Participants had normal hearing, as per a screening pure tone audiogram from 0.25 Hz to 8 kHz and reported no history of neurological or psychiatric disorders and had normal or corrected-to-normal vision.

The experiment simulated a multi-talker scenario (**Figure 1A**) with a circular array (1.50m radius) of six loudspeakers surrounding the listener (at horizontal angles of ±30°, ±112.5°, and ±157.5° relative to the participant). Participants were instructed to dynamically switch their attention between left and right speech streams in the foreground, following a visual cue (left- or right-pointing arrow) indicating the to-be-attended side, which was displayed at the centre of a screen placed in front of them. While switching their attention between the foreground streams, they were also asked to ignore a 16-talker noise played from the four loudspeakers in the background (B1-B4, each of them delivering a 4-talker babble). Frontal streams were presented at 60dB sound pressure level (SPL) each, while each of the noise babbles was delivered at a level of 54dB SPL, resulting in a 3dB signal-to-noise ratio (SNR) of the foreground relative to the background.

Participants were presented with twenty trials (lasting 180 s each) and had to perform 6 attention switches per trial, occurring at semi-random intervals (**Figure 1B**). For this reason, blocks of sustained attention to one particular speech stream varied considerably in duration, spanning from 10s to 30s. For each trial, a different male and female speech stream was played from the left and right loudspeakers in the foreground, counterbalancing throughout the experiment for side of presentation and start of the attention block (i.e., the experiment consisted of five sub-blocks with the following trial sequence: Male Left – Attention Start: Left; Female Left – Attention Start: Left; Male Left – Attention Start: Right; Female Left – Attention Start: Right).

Each trial started with the visual cue pointing towards the to-be-attended side and background noise only, followed by the two foreground speech streams starting simultaneously after 5s. At the end of the trial, participants answered three multiple-choice questions about: the content of the speech streams, their preference based on personal interest (left or right stream), and the perceived difficulty of the attention switching task. The experiment flow was self-paced and, to minimise fatigue, it included three mandatory breaks, each lasting no less than five minutes, every fifth trial.

Speech streams were presented at a sampling rate of 44.1 kHz, delivered through a Roland Octa-Capture 10x10 sound card (24-bit/192 kHz), and played through six PreSonus Eris 4.5BT loudspeakers. Participants’ EEG activity was recorded using a BioSemi ActiveTwo system at a sampling rate of 512 Hz, from sixty-four electrodes positioned on a standard cap following the International 10/20 system. An active (CMS) and a passive (DRL) electrode were used as reference for all electrodes, and two additional electrodes were placed on the mastoids for offline referencing. For 21 of our 24 participants, we additionally recorded electro-oculography (EOG) and electro-myography (EMG). Two electrodes were placed on the left and right temples to capture horizontal eye movements, and two electrodes were positioned above and below the left eye to record vertical eye movements and blinks. To capture EMG activity related to head rotation, an electrode was placed on the left deltoid muscle. Please note that activity from these external electrodes has not been analysed as part of this study.

### Stimuli

The foreground speech stimuli included forty TED Talks covering a range of topics, with 20 female and 20 male presenters, each speaking in a variety of English accents. All speech streams were root-mean-squared (RMS) normalised to reduce differences between male and female voices. Each of the 4-talker background babble signals was obtained by summing the audio signals of four separate TED talks. The long-term average spectrum of the babble noise was then adjusted to align with the overall spectrum of both male and female foreground speakers, to prevent inconsistencies in masking.

### EEG data preprocessing

Neural data were analysed with custom scripts in MATLAB software (MathWorks), based on publicly available scripts and resources shared as part of the CNSP initiative (Cognition and Natural Sensory Processing; https://cnspworkshop.net). Neural signals were first band-pass filtered between 0.5 Hz and 8 Hz, using a zero-phase shift Butterworth filters of order 4, and then downsampled from 512 Hz to 64 Hz. Spherical spline interpolation was applied to replace channels that were three standard deviations away from the mean. EEG was then re-referenced to the average of the two mastoid channels.

### Speech features

The current study aimed to characterise the neural tracking of a dynamic multi-talker scenario by measuring the relationship between EEG data and various features of the foreground speech stimuli, related to their acoustic and lexical properties. To model the speech acoustics, the audios’ broadband amplitude envelopes were extracted by taking the absolute of the Hilbert transform. In order to model the lexical properties of speech, the transcribed stimuli and their audios were first automatically aligned using the WebMAUS Basic aligner ^71–73^, which identified timestamps corresponding to the start and end of each word. The resulting automatic alignment was saved in the TextGrid format and adjusted manually, when necessary, using the Praat software ^74^. The time stamps were then used to build binary word onset vectors in MATLAB. While binary word onset vectors represent information related to word segmentation, they can also be modulated according to each word’s surprisal or entropy value to represent higher-order semantic information. Word surprisal is a measure of how unexpected a word is given its preceding linguistic context, and it can be computed using Large Language Models (LLMs), as the negative logarithm of the probability of that word given the previous context. Word entropy, on the other hand, measures how uncertain or unpredictable the next word is. Here, we used a pretrained open-source LLM, Mistral-7B-v0.1 ^75^ to extract word probabilities, and then computed lexical surprisal and lexical entropy values for each word.

When considering the dynamic nature of the attention switching task, the definition of what context to include for the current word prediction becomes a non-trivial problem. From the machine perspective, given a word *w* belonging to, e.g., the left stream, the LLM would predict it more easily when provided with all the available linguistic context from the left stream. However, from the neural/behavioural perspective, since participants were flexibly re-orienting their attention between left and right streams, the optimal context would be impacted by the attention switch and, potentially, store information of previously attended blocks. To compare these context-accumulation alternatives, we represented context according to four alternative representations. A machine-ideal model, Oracle, considers as context all words preceding the current word in one particular stream, whether they were attended or unattended. As such, this model is *unaware* of the switch of attention. Among more neurally-plausible and switch-aware context accumulation models, we constructed the Attention model, which incorporates as context any previously attended block from both left and right speech streams, and the Speaker Specific model, which displays a speech stream bias, whereby only previously attended blocks of the same stream are included as context for lexical prediction of the current block. Similarly to Attention and Speaker Specific, the fourth model, Reset, is switch-aware, but not context-aware. In fact, it does not keep track of any previous speech block, neither attended nor unattended, and instead resets the context following each attention switch cue.

### Temporal response function (TRF) and analysis procedure

In this study, we quantified neural tracking of speech features as the ability to predict unseen EEG data (encoding) or reconstruct unseen stimulus features (decoding), with a linear model based on lagged linear regression, the Temporal Response Function (TRF). A TRF model was fit for each participant separately, using a leave-one-trial-out cross-validation procedure to avoid overfitting, and considering a range of time lags from -100ms to 600ms in the case of the encoding models, and between 0ms and 600ms for the decoding models. The TRF model calculation applied Tikhonov regularisation, with the regularisation parameter (λ) tuned via an exhaustive search across a logarithmic scale from 10^-6^ to 10^4^. This search was conducted on the training fold during each iteration of the cross-validation process, and the optimal λ was selected as the parameter that yielded the highest correlation (Pearson’s *r*) between true and predicted/reconstructed data.

We first constructed an Attended speech model from the combination of attended segments of left and right speech streams. We then fitted TRF models by coupling the EEG and the Attended stimulus features. With a decoding approach, we reconstructed the envelope of each trial based on the Attended TRF model, and then correlated it with the true envelopes of left and right streams for the corresponding trial. Since the left or right true attention label was known throughout the trial, classification accuracy could be computed as the percentage of left or right segments correctly classified as attended (**Figure 1C**). We also computed the null distribution of the classification accuracy by randomly shuffling the left and right labels for 100 iterations. We employed a range of decoding window lengths (1 s,2 s,4 s,8 s,16 s,32 s) to assess the impact of correlation window size on the model’s ability to correctly classify the attended stimulus.

To observe the evolution of neural tracking of left and right streams over time with a neurophysiologically interpretable approach, we calculated an encoding TRF model where true and predicted EEG signals were correlated using a sliding window approach (MATLAB’s *movcorr* function) with a step size of 0.125s. The encoding TRFs were fit with a multivariate set of attended stimulus features including envelope (Env), word onset (WO) and word surprisal (WS) or word entropy (WE), to assess the joint impact of both acoustic and semantic features on the temporal dynamics of the attention switch.

The re-orienting of attention around the switch can be analysed in terms of disengaging stream (attended > unattended) and engaging (unattended > attended) stream. To compare the onset and offset points of these processes, we first selected participants depending on their attentional bias over the course of the switch. Based on the reference 4-s encoding window, we determined the single-subject classification accuracy in a 15s window before and after the attention switching cue. We then found those participants displaying a lower-than-chance (50%) classification performance either in the pre- or post-switch window and excluded them from further analyses on the switching temporal dynamics. Start and end points of disengagement and engagement processes were identified for each subject by fitting a piecewise linear regression model to the EEG prediction correlation values for the disengaging and the engaging streams, around the attention switch cue. For each of the EEG prediction correlation streams, the model incorporates the average pre-switch and post-switch values as constants and tries to estimate the breakpoints at which these flat segments connect, i.e. the onset and offset points of the process. An initial guess for the breakpoints was provided based on reasonable assumptions (e.g., that the start of the disengagement/engagement could not occur before the switching cue), and the fitting was done with MATLAB’s *lsqcurvefit* function, which minimises the difference between the model and the observed data.

When computing encoding TRFs comparing different kinds of context-accumulation models, we adopted an approach optimised for highlighting the effects of lexical information processing ^76^. With this method, the prediction correlation between true and predicted EEG data is computed within a window covering the range from 300ms to 500ms after word onset, allowing to effectively enhance the impact of semantic encoding.

### Event-related spectral perturbation (ERSP)

Previous studies have related decreased parietal alpha-band (8-12 Hz) EEG power to increased listening effort and have shown that more listening effort is required to switch attention than to sustain attention on a particular speaker ^29^. Since we were interested in assessing the time course of alpha power and how it would compare to the temporal dynamics of the TRF prediction correlations, we computed event-related spectral perturbation (ERSP) in the alpha frequency band as a proxy of effort around the switch. Since neural activity in the alpha frequency band is sensitive to ocular artifacts, we performed Independent Component Analysis (ICA) using standard EEGLAB functions ^77^. For each subject, we identified independent components reflecting ocular artifacts (such as blinks and horizontal eye movements) based on their temporal characteristics and spatial topographies, and we removed them from the data. In this study, ERSP is a measure of the alpha power as a change relative to a 15s pre-switch baseline. Power spectral density (PSD) was estimated via Welch’s method (MATLAB’s *pwelch* function) for each trial, channel, and switch. For this analysis, EEG data was filtered between 0.5 Hz and 20 Hz, but only PSD values between 8 Hz and 12 Hz frequency bands were included to compute the absolute alpha power (squared sum). While the baseline PSD included the whole 15s of pre-switch window, the PSD following the attention switching cue was computed with a sliding window of 1,2,4 and 8 seconds, in order to compare it directly to the TRF correlations. For each of the windows, the relative alpha power was calculated as the percentage difference compared to the baseline PSD.

### Statistical analysis

Statistical analyses were performed using repeated measures ANOVAs with two or three factors, with *F*-values reported as *F*(df_time_, df_error_). When the assumption of sphericity wasn’t met, as revealed by Mauchly’s test of sphericity, a correction was applied with the Greenhouse-Geisser’s method. When a significant main effect was found, post-hoc tests were run, and *p*-values were corrected for multiple comparisons with the Holm method. To compare the significance of a time series against zero, a Wilcoxon signed rank test was used for each time point, and the false discovery rate (FDR) was controlled with the Benjamini & Hochberg procedure. We report FDR-adjusted *p*-values. Descriptive statistics for the neurophysiology results are reported as a combination of mean and standard error (SE).

## Acknowledgements

This work was conducted with the financial support of the William Demant Fonden (https://www.williamdemantfonden.dk/), grant 21-0628 and grant 22-0552 and of Taighde Éireann – Research Ireland (https://www.researchireland.ie/), under Grant No. 18/CRT/6223. This research was supported by the Research Ireland under Grant Agreement No. 13/RC/2106_P2 at the ADAPT SFI Research Centre (https://www.sfi.ie/sfi-research-centres/adapt/) at Trinity College Dublin. ADAPT, the Research Ireland Centre for AI-Driven Digital Content Technology, is funded by Research Ireland through the RI Research Centres Programme.

