## Supplemental Figure 1 for "Simultaneous cortical tracking of competing speech streams during attention switching"

[7] Senior author.

[8] Lead contact.

### Supplemental information

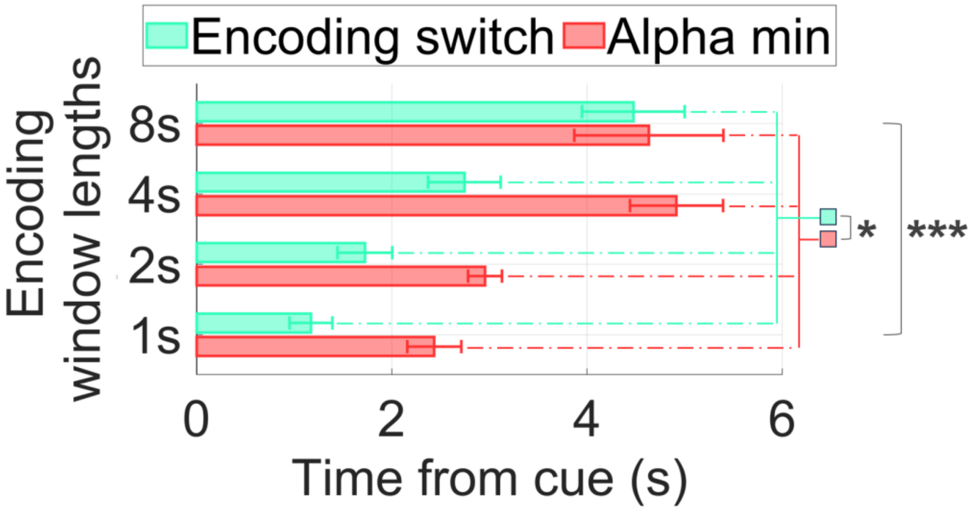


**Figure S1. Related to Figure 2.** Time latencies of the encoding switch of EEG prediction correlations (turquoise) and alpha ERSP minimum (red) for four sliding window lengths. Bars represent the average time latencies across participants and electrodes, error bars indicate the SEM across participants. Stars represent statistically significant comparisons (Significance levels: *$p < 0.05$, **$p < 0.01$, ***$p < 0.001$).
